# Gangliosides in neural stem cell fate determination and nerve cell specification--preparation and administration

**DOI:** 10.1101/2024.06.09.598109

**Authors:** Yutaka Itokazu, Toshio Ariga, Takahiro Fuchigami, Dongpei Li

**Author notes:** Y.I. and T.A. contributed equally to this work. **Correspondence:** Yutaka Itokazu.

## Abstract

Gangliosides are sialylated glycosphingolipids with essential but enigmatic functions in healthy and disease brains. GD3 is the predominant species in neural stem cells (NSCs) and GD3-synthase (sialyltransferase II; *St8Sia1*) knockout (GD3S-KO) revealed reduction of postnatal NSC pools with severe behavioral deficits including cognitive impairment, depression-like phenotypes, and olfactory dysfunction. Exogenous administration of GD3 significantly restored the NSC pools and enhanced the stemness of NSCs with multipotency and self-renewal, followed by restored neuronal functions. Our group discovered that GD3 is involved in the maintenance of NSC fate determination by interacting with epidermal growth factor receptors (EGFRs), by modulating expression of cyclin-dependent kinase (CDK) inhibitors p27 and p21, and by regulating mitochondrial dynamics via associating a mitochondrial fission protein, the dynamin-related protein-1 (Drp1). Furthermore, we discovered that nuclear GM1 promotes neuronal differentiation by an epigenetic regulatory mechanism. GM1 binds with acetylated histones on the promoter of *N-acetylgalactosaminyltransferase (GalNAcT; GM2 synthase (GM2S); B4galnt1)* as well as on the *NeuroD1* in differentiated neurons. In addition, epigenetic activation of the GM2S gene was detected as accompanied by an apparent induction of neuronal differentiation in NSCs responding to an exogenous supplement of GM1. Interestingly, GM1 induced epigenetic activation of the *tyrosine hydroxylase (TH)* gene, with recruitment of Nurr1 and PITX3, dopaminergic neuron-associated transcription factors, to the *TH* promoter region. In this way, GM1 epigenetically regulates dopaminergic neuron specific gene expression, and it would modify Parkinson’s disease. Multifunctional gangliosides significantly modulate lipid microdomains to regulate functions of important molecules on multiple sites: the plasma membrane, mitochondrial membrane, and nuclear membrane. Versatile gangliosides regulate functional neurons as well as sustain NSC functions via modulating protein and gene activities on ganglioside microdomains. Maintaining proper ganglioside microdomains benefits healthy neuronal development and millions of senior citizens with neurodegenerative diseases.

Here, we introduce how to isolate GD3 and GM1 and how to administer them into the mouse brain to investigate their functions on NSC fate determination and nerve cell specification.

## 1. Introduction

Glycosphingolipid composition, including gangliosides [1], is dynamically changed during neural differentiation (GD3 => GM1), thereby regulating distinct stages of cell fate determination. Our and others’ studies suggest that GD3 maintains neural stem cell (NSC) characteristics and that GM1 epigenetically promotes neuronal differentiation and maintains neuronal functions [2-17] (Fig. 1).

**Fig. 1.**
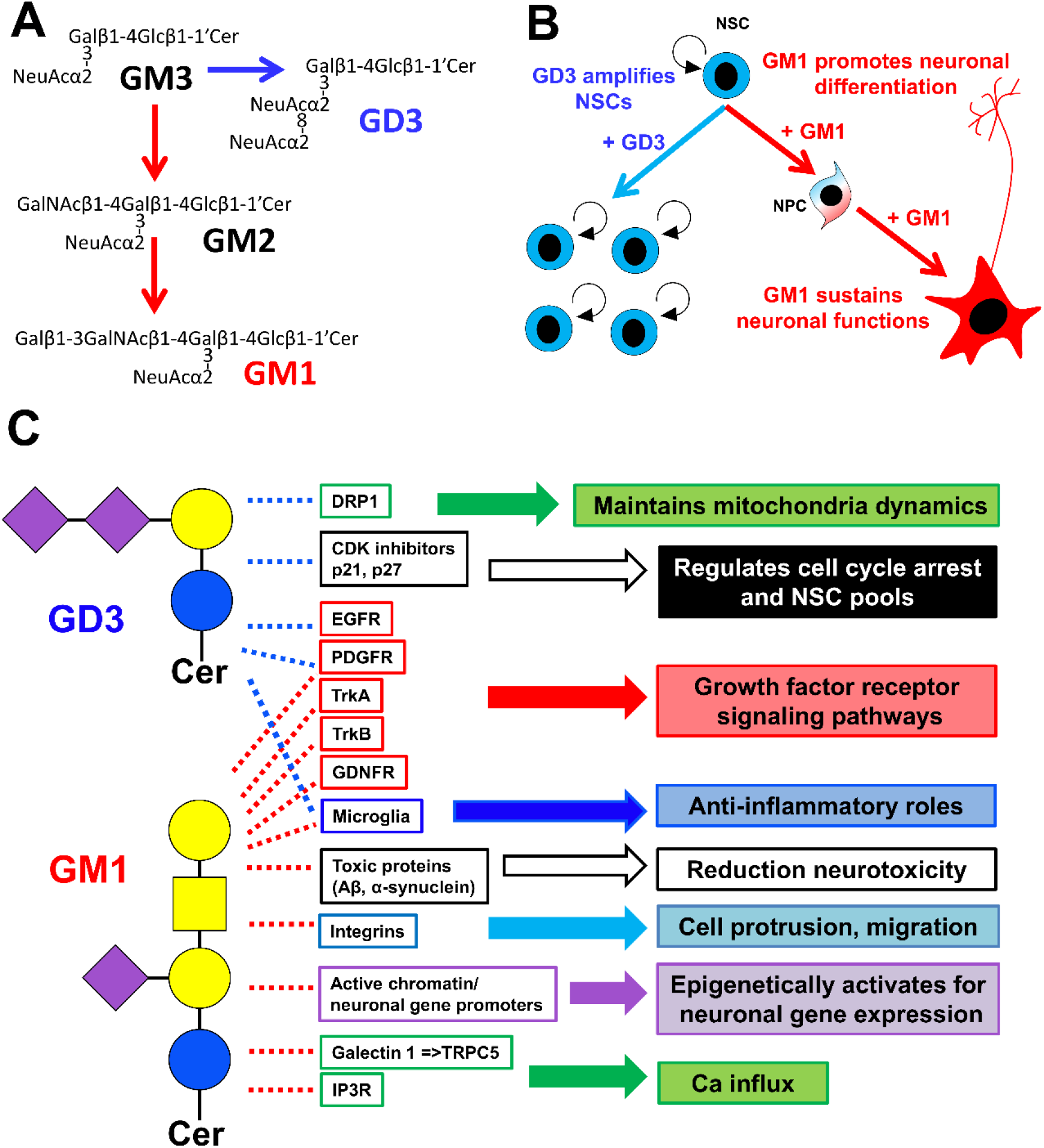
Metabolic pathways and structure of GD3 and GM1 gangliosides. (A) The nomenclature for gangliosides and their components are based on that of Svennerholm and the IUPAC–IUBMB Joint Commission on Biochemical Nomenclature [18, 19]. (B) GD3 restores the number of neural stem cells (NSCs), and GM1 restores neuronal precursor cells (NPCs) and neuronal functions. (C) Examples of multifunction of GD3 and GM1: Without GD3, increased Drp1 levels induce aberrant mitochondrial fragmentation. GD3 suppresses p21 expression and maintains p27 expression in NSCs. Gangliosides bind to the neurotrophic factor receptors to regulate their signaling. Gangliosides decrease inflammatory microglia & increase phagocytosis. Gangliosides prevent and even reduce the accumulation of toxic proteins [e.g., amyloid-beta peptides (Aβs) and alpha-synuclein (α-syn)] in neurodegenerative diseases. GM1 promotes integrin signaling for cell protrusion including neurite outgrowth and cell migration. Nuclear GM1 epigenetically promotes neuronal gene expression to sustain healthy neuronal functions. GM1 induces Ca influx.

Functional roles of gangliosides have been studied with ganglioside knockout (KO) mice (i.e., glycosyltransferase KO mice). However, KO mice cannot exclude the effect of ganglioside metabolic changes (e.g., upregulated, or downregulated other than interesting molecules). To elucidate the functional roles of gangliosides, exogenous administration of ganglioside(s) is a critical and desirable experiment. In previous studies, intracerebroventricular administration was the most reliable method of delivering gangliosides into the brain [8, 20, 21]. We have successfully developed a more convenient non-invasive delivery procedure to the brain by intranasal infusion of gangliosides [3, 7, 22, 23]. In the present work, we review the laboratory methods that one can use to study the ganglioside function on NSC fate determinations in the mouse brain.

## 2. Materials

1. Chloroform.
2. Methanol.
3. Sodium acetate.
4. Ammonia.
5. n-propanol (1-propanol).
6. Calcium chloride.
7. DEAE-Sephadex A-25.
8. Iatrobeads.
9. Glass chromatography columns (5 cm inner diameter (i.d.) X 150 cm; 2.5 cm i.d. X 200 cm; 2.2 cm i.d. X 100 cm; 1.5 cm i.d. X 100 cm; 1.2 cm i.d. X 56 cm; 2 cm i.d. X 100 cm; 0.4 cm i.d. X 110 cm).
10. Bovine, porcine, ovine, or human brains (for GM1). Bovine buttermilk powder (for GD3).
11. Saline (0.9% sodium chloride).
12. Anesthetic agent (e.g., ketamine hydrochloride and xylazine hydrochloride).
13. Stereotaxic apparatus.
14. Mini-osmotic pumps (e.g., ALZET, model #1007D, 100 μl capacity for 7-day infusion).
15. Capillary tips (Bio-Rad, #2239915).

## 3. Methods

### 3.1. Purification of GM1

The procedure of GM1 isolation is based on Ando and Yu [24] with minor modifications of Ariga and Yu [25].

1. Homogenize the brain (about 2.3 kg) with ten volumes each of chloroform/methanol 2:1, 1:1, and 1:2 (v/v).
2. Combine all lipid extracts.
3. Evaporate solvent.
4. Dissolve the residue in12 liters of chloroform/methanol/water 30:60:8 (v/v/v) (Solvent A).
5. Apply to a DEAE-Sephadex A-25 column chromatography (acetate form, 5 cm i.d. X 150 cm, 310 g).
6. Elute the neutral lipids with fifteen liters of chloroform/methanol/water 30:60:8 (v/v/v) and two liters of methanol.
7. Add thirteen liters of 0.2 M sodium acetate in methanol to recover the acidic lipids.
8. Evaporate solvent.
9. Dissolve the residue in one liter of water.
10. Dialyze against distilled water for 3 days.
11. Lyophilize the retentate.
12. Dissolve the residue in 500 ml of chloroform/methanol/water 30:60:8 (v/v/v)
13. Apply to a DEAE-Sephadex A-25 column chromatography (2.5 cm i.d. X 200 cm).
14. At this stage, there are still lesser amounts of residual neutral lipids. Elute the neutral lipids with1 liter of chloroform/methanol/water 30:60:8 (v/v/v) and 500 ml of methanol.
15. Elute the acidic glycolipids with four liters of a linear gradient system prepared from sodium acetate in methanol (0.05 M and 0.3 M).
16. Combine the monosialoganglioside fractions.
17. Evaporate solvent.
18. Dissolve the residue in 200 ml of distilled water.
19. Dialyze against distilled water.
20. Lyophilize the retentate.
21. Dissolve the residue in 20 ml of chloroform/methanol/water 70:30:1 (v/v/v).
22. Apply to an Iatrobeads column chromatography (2.2 cm i.d. X 100 cm).
23. Elute from column with two liters of a linear gradient system of chloroform/methanol/water 65:35:4 and 45:55:5 (v/v/v).
24. Combine the fraction containing GM1.
25. Evaporate solvent.
26. Dissolve the residue in 20 ml of chloroform/methanol/water 70:30:1 (v/v/v).
27. Apply again to another Iatrobeads column chromatography (1.5 cm i.d. X 100 cm).
28. Elute from column with one liter of a linear gradient system of chloroform/methanol/water 65:35:4 and 45:55:5 (v/v/v).
29. Evaporate solvent.
30. Dissolve the residue in 20 ml of chloroform/methanol/water 70:30:1 (v/v/v).
31. Apply the final Iatrobeads column chromatography (1.2 cm i.d. X 56 cm).
32. Elute from column with 500 ml of n-propanol/water 85:15 (v/v).
33. Confirm the purity of GM1 by high-performance thin-layer chromatography (HPTLC) with the following solvent system: (a) chloroform/methanol/0.02% calcium chloride 50:45:10 (v/v/v) and (b) chloroform-methanol/5M ammonia/0.4% calcium chloride 60:45:4:5 (v/v/v); and/or by Mass spectrometry (MS)-based ganglioside profiling.

### 3.2. Purification of GD3

The procedure of GD3 isolation is based on previous publications [26, 27].

1. Suspend the bovine buttermilk powder (1.7 kg, containing 32% protein, 8.5% mineral, 49% carbohydrates, and 3.5% other small molecular weight compounds) in water.
2. Dialyze against distilled water for 3 days to remove small oligosaccharides, minerals, and other small molecular weight materials.
3. Lyophilize the retentate.
4. Extract total lipids with ten volumes each of chloroform/methanol 2:1, 1:1, and 1:2 (v/v).
5. Combine all lipid extracts.
6. Adjust to a ratio of chloroform/methanol/water 2:1:0.6 (v/v/v).
7. Partition three times according to the procedure of Folch et al. (1957) [28].
8. Combine the ganglioside fractions from the upper phase.
9. Evaporate solvent.
10. Dialyze the residue against distilled water for 3 days with frequent changing of water.
11. Lyophilize the retentate.
12. Adjust the residue with chloroform/methanol/water 30:60:8 (v/v/v).
13. Apply to a DEAE-Sephadex A-25 column chromatography (2 cm i.d. X 100 cm).
14. Remove neutral lipids with ten volumes of chloroform/methanol/water 30:60:8 (v/v/v).
15. Elute the crude gangliosides by stepwise elution with a solvent system of one liter each of 0.05 M, 0.2 M, 0.8 M sodium acetate in chloroform/methanol-/water 30:60:8 (v/v/v).
16. Collect each 15-ml of effluent fractions.
17. Pool the fractions 1-80 as monosialogangliosides, 80-110 as disialogangliosides, and 110-140 as trisialo-gangliosides.
18. Evaporate solvent.
19. Dissolve the residue in 20 ml of chloroform/methanol/water 70:30:1 (v/v/v).
20. Apply to an Iatrobeads column chromatography (2 cm i.d. X 100 cm).
21. Elute from column with two liters of a linear gradient elution using one liter each of chloroform/methanol/water 70:30:3 and 35:65:5 (v/v/v) to yield GD3.
22. Evaporate solvent.
23. Dissolve the residue in 20 ml of chloroform/methanol/water 70:30:1 (v/v/v).
24. Apply the final Iatrobeads column chromatography (0.4 cm i.d. X 110 cm).
25. Elute from column with one liter of a linear gradient elution using one liter each of chloroform/methanol/water 70:30:3 and 35:65:5 (v/v/v) to yield GD3.
26. Confirm the purity of GD3 by high-performance thin-layer chromatography (HPTLC) with the following solvent system: (a) n-propanol/water 80:20 (v/v), (b) chloroform/methanol/0.5 % calcium chloride 55:45:10 (v/v/v); and/or by Mass spectrometry (MS)-based ganglioside profiling.

### 3.3. Intracerebroventricular administration of gangliosides

The procedure of intracerebroventricular administration of gangliosides (Fig. 2) is based on Itokazu et al. [8].

**Fig. 2.**
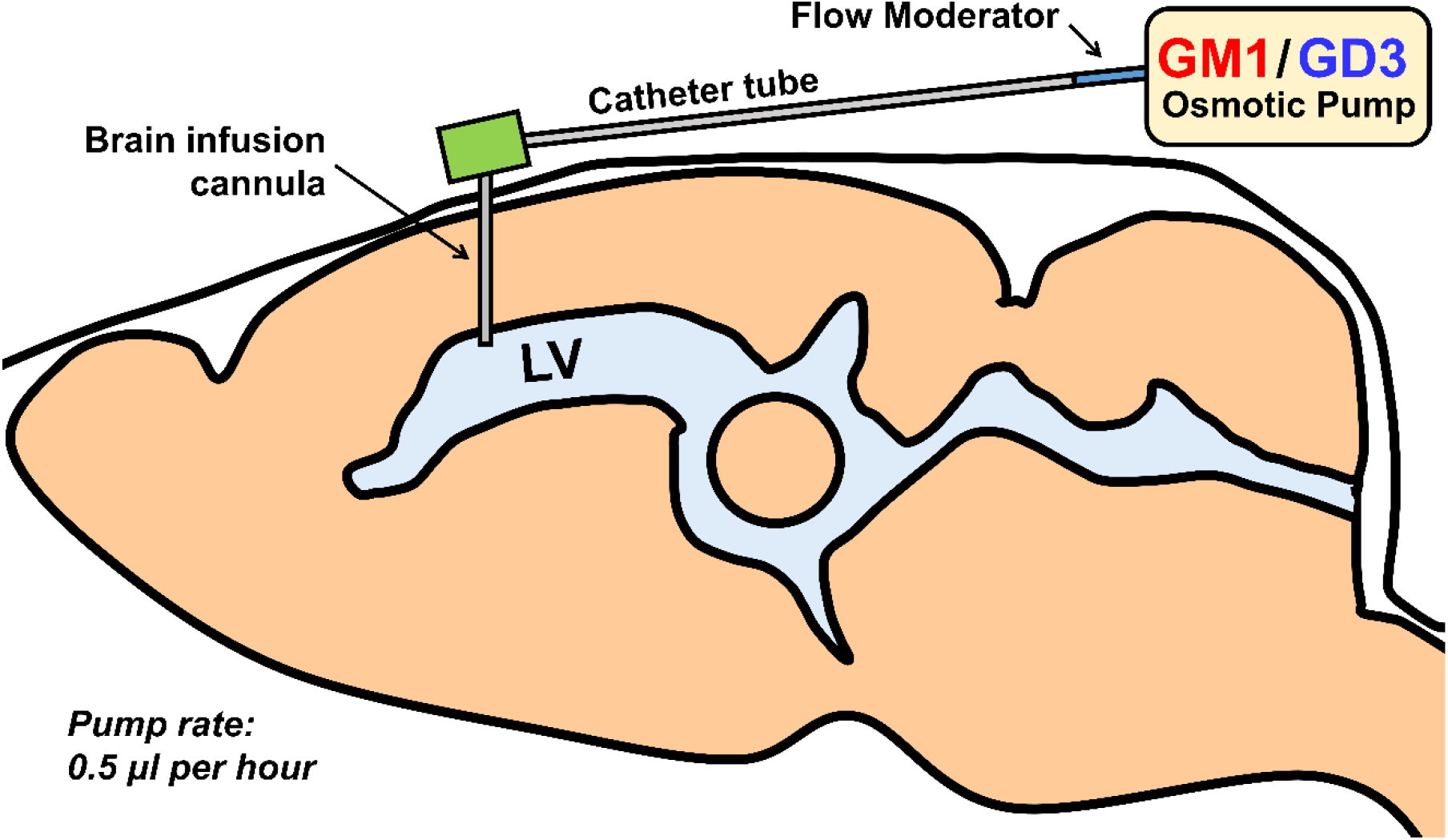
Intracerebroventricular administration of gangliosides using mini-osmotic pump. Implant the brain infusion cannula into the lateral ventricle, and the pump (filled with GD3 or GM1) attached by tubing to the brain infusion cannula subcutaneously in the neck region of the back.

1. Prepare ganglioside solution in saline (5 mg/kg-weight/day, twelve μl for a daily infusion).
2. Load ganglioside solution into the Mini-osmotic pumps (ALZET, Cupertino, CA) and connect the catheter tube and cannula as instructed on the manual.
3. Anesthetize the mouse with ketamine hydrochloride (40 mg/kg body weight) and xylazine hydrochloride (4 mg/kg body weight).
4. Fix the mouse on a stereotaxic apparatus.
5. Shave and disinfect the scalp; make a midline sagittal incision and expose the skull; and prepare a subcutaneous pocket in the back of the mice to hold the osmotic pump.
6. Identify the bregma as the reference point. Move the tip of the stereotaxic coordinates to AP: 0.4 mm, ML: 1.5 mm. Drill a hole in the skull. Adjust the cannula to DV: 3 mm and apply glue to the cannula holder. Inset the cannula into the desired place and direct pressure should be applied to make sure the cannula holder is attached to the skull tightly.
7. Place the osmotic pump in the subcutaneous pocket.
8. Close the scalp and the skin of the back with suture or skin glue.

### 3.4. Intranasal administration of gangliosides

The procedure of intranasal administration of gangliosides (Fig. 3) is based on Itokazu et al. [7] and Fuchigami et al. [3].

**Fig. 3.**
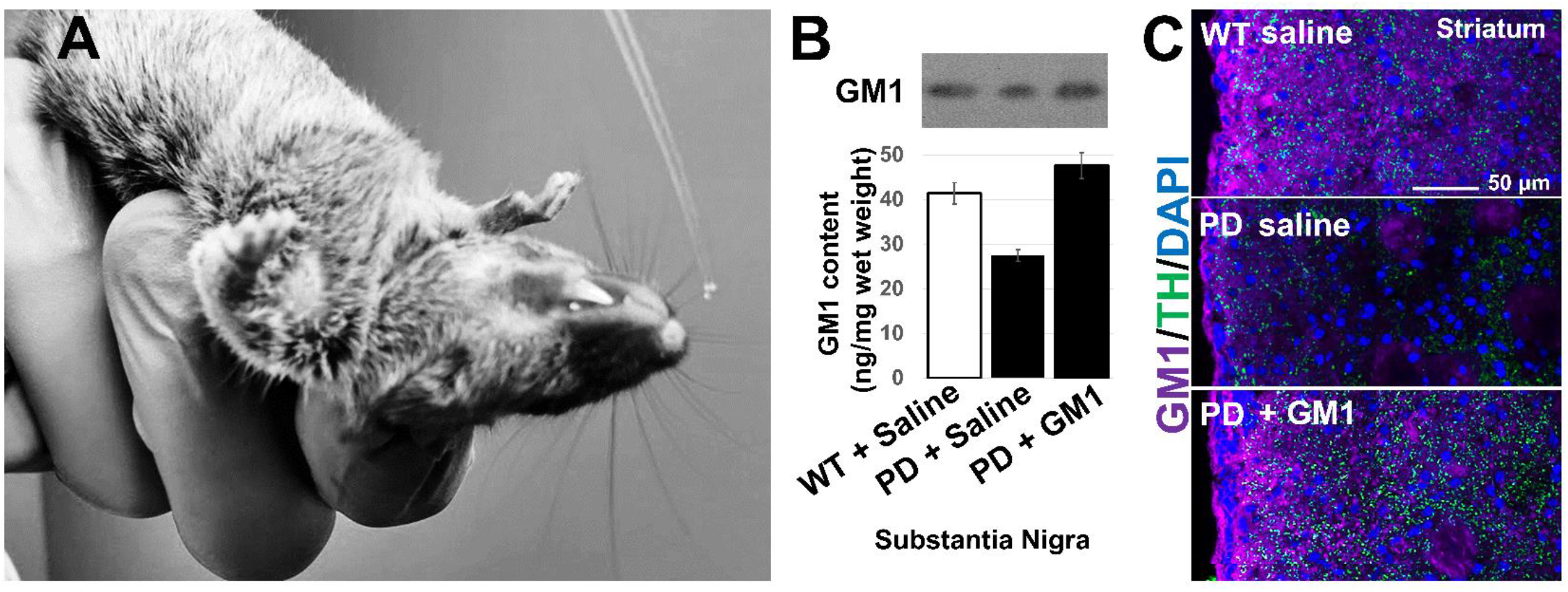
Intranasal administration of gangliosides. (A) Intranasal drug delivery is an alternative administration, using the direct anatomic pathway between the olfactory/trigeminal neuroepithelium of the nasal mucosa and the brain. (B) Nasal GM1 (5 mg/kg/day for 28 days) restored GM1 levels in the Parkinson’s disease (PD) model (A53T) mouse brain (by HPTLC). (C) Intranasally infused GM1 (5 mg/kg/day for 28 days) recovered the tyrosine hydroxylase (TH)+ axons (green) in the striatum of PD model mice. This suggests that chronic dopamine depletion seen in the PD-model mice and possibly of PD patients might be restored by intranasal GM1.

1. Prepare ganglioside solution in saline (5 mg/kg-weight/day, twenty-four μl for a daily infusion).
2. Hold the mouse on the back of the neck using the thumb and pointer finger.
3. Invert with the ventral side facing up towards the ceiling.
4. Infuse six μl of ganglioside solution or saline into the naris of the mouse using capillary tips (Fig. 3A).
5. Keep holding the mouse in this position for 30 seconds.
6. Repeat the procedure for an additional three times. Infuse each six μL of ganglioside solution into the right and left nares twice (total twenty-four μl infusion).
7. Intranasally infuse daily for desired days.

## 4. Notes

1. Momoi et al. [29] reported the purification of large amount of GM1 from bovine brain. Sugimoto et al., reported the chemical synthesis of GM1 [30].
2. Chemical synthesis of GD3 was reported by Ito et.al. [31] and Ishida et.al. [32].
3. Purified lipids can be stored in the freezer.
4. All animal experiments were approved by the Institutional Animal Care and Use Committee (IACUC) at Augusta University (AU) according to the National Institutes of Health (NIH) guidelines and were performed with approved animal protocols (references AUP 2009-0240 and 2014-0694).
5. Ganglioside treated mice will be sacrificed for further NSC analyses.
6. In our experiments, no detectable adverse effects occurred during or after intranasal administration of gangliosides. Tissue sections were analyzed with hematoxylin and eosin (HE) staining, and no abnormal structures were found in the olfactory epithelium, brain, eye, spinal cord, tongue, thymus, lung, liver, spleen, kidney, stomach, intestine, colon, bone, or bone marrow after intranasal GD3 or GM1 infusion. Neither histological damage nor tumor formation was observed. Enzyme-linked immunosorbent assay (ELISA) revealed that intranasal ganglioside infusion (28 days) did not elevate the titer of ganglioside antibodies in the serum. Thus, intranasally administered gangliosides are safe and show no toxicity.

## Acknowledgments

This work was partly supported by a National Institute of Neurological Disorders and Stroke grant (Y.I.: R01NS100839), a Sheffield Memorial Grant of the Central Savannah River Area (CSRA) Parkinson Support Group (Y.I.), and the excellent infrastructural support of the Department of Pharmacology and Toxicology (Chair: Dr. Alvin Terry), Medical College of Georgia at Augusta University.

## References

1. Yu RK, Tsai YT, Ariga T and Yanagisawa M (2011) Structures, biosynthesis, and functions of gangliosides--an overview. Journal of oleo science 60(10):537–544

2. Fantini J and Barrantes FJ (2009) Sphingolipid/cholesterol regulation of neurotransmitter receptor conformation and function. Biochimica et biophysica acta 1788(11):2345–2361 doi:10.1016/j.bbamem.2009.08.016

3. Fuchigami T, Itokazu Y, Morgan JC and Yu RK (2023) Restoration of Adult Neurogenesis by Intranasal Administration of Gangliosides GD3 and GM1 in The Olfactory Bulb of A53T Alpha-Synuclein-Expressing Parkinson’s-Disease Model Mice. Molecular neurobiology doi:10.1007/s12035-023-03282-2

4. Fuchigami T, Itokazu Y and Yu RK (2024) Ganglioside GD3 regulates neural stem cell quiescence and controls postnatal neurogenesis. Glia 72(1):167–183 doi:10.1002/glia.24468

5. Furukawa K et al. (2019) New era of research on cancer-associated glycosphingolipids. Cancer Sci 110(5):1544–1551 doi:10.1111/cas.14005

6. Galleguillos D et al. (2022) Anti-inflammatory role of GM1 and other gangliosides on microglia. J Neuroinflammation 19(1):9 doi:10.1186/s12974-021-02374-x

7. Itokazu Y, Fuchigami T, Morgan JC and Yu RK (2021) Intranasal infusion of GD3 and GM1 gangliosides downregulates alpha-synuclein and controls tyrosine hydroxylase gene in a PD model mouse. Mol Ther 29(10):3059–3071 doi:10.1016/j.ymthe.2021.06.005

8. Itokazu Y, Li D and Yu RK (2019) Intracerebroventricular Infusion of Gangliosides Augments the Adult Neural Stem Cell Pool in Mouse Brain. ASN neuro 11:1759091419884859 doi:10.1177/1759091419884859

9. Itokazu Y, Tsai YT and Yu RK (2016) Epigenetic regulation of ganglioside expression in neural stem cells and neuronal cells. Glycoconjugate journal doi:10.1007/s10719-016-9719-6

10. Itokazu Y, Wang J and Yu RK (2018) Gangliosides in Nerve Cell Specification. Progress in molecular biology and translational science 156:241–263 doi:10.1016/bs.pmbts.2017.12.008

11. Ledeen RW and Wu G (2015) The multi-tasked life of GM1 ganglioside, a true factotum of nature. Trends in biochemical sciences 40(7):407–418 doi:10.1016/j.tibs.2015.04.005

12. Schneider JS, Singh G, Williams CK and Singh V (2022) GM1 ganglioside modifies microglial and neuroinflammatory responses to alpha-synuclein in the rat AAV-A53T alpha-synuclein model of Parkinson’s disease. Molecular and cellular neurosciences 120:103729 doi:10.1016/j.mcn.2022.103729

13. Tang FL, Wang J, Itokazu Y and Yu RK (2021) Ganglioside GD3 regulates dendritic growth in newborn neurons in adult mouse hippocampus via modulation of mitochondrial dynamics. J Neurochem 156(6):819–833 doi:10.1111/jnc.15137

14. Tsai YT, Itokazu Y and Yu RK (2016) GM1 Ganglioside is Involved in Epigenetic Activation Loci of Neuronal Cells. Neurochemical research 41(1-2):107–115 doi:10.1007/s11064-015-1742-7

15. Tsai YT and Yu RK (2014) Epigenetic activation of mouse ganglioside synthase genes: implications for neurogenesis. J Neurochem 128(1):101–110 doi:10.1111/jnc.12456

16. Wang J, Cheng A, Wakade C and Yu RK (2014) Ganglioside GD3 is required for neurogenesis and long-term maintenance of neural stem cells in the postnatal mouse brain. The Journal of neuroscience : the official journal of the Society for Neuroscience 34(41):13790–13800 doi:10.1523/JNEUROSCI.2275-14.2014

17. Wang J and Yu RK (2013) Interaction of ganglioside GD3 with an EGF receptor sustains the self-renewal ability of mouse neural stem cells in vitro. Proceedings of the National Academy of Sciences of the United States of America 110(47):19137–19142 doi:10.1073/pnas.1307224110

18. Svennerholm L (1963) Chromatographic Separation of Human Brain Gangliosides. J Neurochem 10:613–623

19. [No authors listed] (1977) The nomenclature of lipids. (Recommendations 1976) IUPAC-IUB Commission on Biochemical Nomenclature. Lipids 12(6):455–468

20. Di Pardo A et al. (2012) Ganglioside GM1 induces phosphorylation of mutant huntingtin and restores normal motor behavior in Huntington disease mice. Proceedings of the National Academy of Sciences of the United States of America 109(9):3528–3533 doi:10.1073/pnas.1114502109

21. Svennerholm L et al. (2002) Alzheimer disease - effect of continuous intracerebroventricular treatment with GM1 ganglioside and a systematic activation programme. Dement Geriatr Cogn Disord 14(3):128–136

22. Itokazu Y, Fuchigami T and Yu RK (2023) Functional Impairment of the Nervous System with Glycolipid Deficiencies. Advances in neurobiology 29:419–448 doi:10.1007/978-3-031-12390-0_14

23. Itokazu Y (2024) Multifunctional glycolipids as multi-targeting therapeutics for neural regeneration. Neural Regen Res 19(4):707–708 doi:10.4103/1673-5374.382244

24. Ando S and Yu RK (1977) Isolation and characterization of a novel trisialoganglioside, GT1a, from human brain. The Journal of biological chemistry 252(18):6247–6250

25. Ariga T and Yu RK (1987) Isolation and characterization of ganglioside GM1b from normal human brain. Journal of lipid research 28(3):285–291

26. Ledeen RW and Yu RK (1982) Gangliosides: structure, isolation, and analysis. Methods Enzymol 83:139–191

27. Ren S et al. (1992) O-acetylated gangliosides in bovine buttermilk. Characterization of 7-O-acetyl, 9-O-acetyl, and 7,9-di-O-acetyl GD3. The Journal of biological chemistry 267(18):12632–12638

28. Folch J, Lees M and Sloane Stanley GH (1957) A simple method for the isolation and purification of total lipides from animal tissues. The Journal of biological chemistry 226(1):497–509

29. Momoi T, Ando S and Magai Y (1976) High resolution preparative column chromatographic system for gangliosides using DEAE-Sephadex and a new porus silica, Iatrobeads. Biochimica et biophysica acta 441(3):488–497

30. Sugimoto M et al. (1986) Total synthesis of gangliosides GM1 and GM2. Carbohydr Res 156:C1–5 doi:10.1016/s0008-6215(00)90125-3

31. Ito Y, Numata M, Sugimoto M and Ogawa T (1989) Highly Stereoselective Synthesis of Ganglioside Gd3 .65. J Am Chem Soc 111(22):8508–8510 doi:DOI 10.1021/ja00204a028

32. Ishida H et al. (1993) A synthetic approach to polysialogangliosides containing alpha-sialyl-(2-->8)-sialic acid: total synthesis of ganglioside GD3. Carbohydr Res 246:75–88 doi:10.1016/0008-6215(93)84025-2

